# The fungal collaboration gradient dominates the root economics space in plants

**DOI:** 10.1101/2020.01.17.908905

**Authors:** Joana Bergmann, Alexandra Weigelt, Fons van der Plas, Daniel C. Laughlin, Thom W. Kuyper, Nathaly Guerrero-Ramirez, Oscar J. Valverde-Barrantes, Helge Bruelheide, Grégoire T. Freschet, Colleen M. Iversen, Jens Kattge, M. Luke McCormack, Ina C. Meier, Matthias C. Rillig, Catherine Roumet, Marina Semchenko, Christopher J. Sweeney, Jasper van Ruijven, Larry M. York, Liesje Mommer

**Affiliations:** Freie Universität Berlin, Germany; Berlin-Brandenburg Institute of Advanced Biodiversity Research (BBIB), Germany; Systematic Botany and Functional Biodiversity, Institute of Biology, Leipzig University, Germany; German Centre for Integrative Biodiversity Research (iDiv) Halle-Jena-Leipzig, Germany; University of Wyoming, USA; Wageningen University, The Netherlands; Biodiversity, Macroecology & Biogeography, Faculty of Forest Sciences and Forest Ecology, University of Göttingen, Germany; International Center of Tropical Botany, Florida International University, USA; Martin Luther University Halle-Wittenberg, Institute of Biology/Geobotany and Botanical Garden, Germany; CEFE, CNRS, Université de Montpellier, Université Paul Valéry Montpellier 3, EPHE, IRD, Montpellier, France; Station d’Ecologie Théorique et Expérimentale (CNRS, Université Toulouse III), Moulis, France; Oak Ridge National Laboratory, USA; MPI Biogeochemistry, Germany; Center for Tree Science, The Morton Arboretum, USA; University of Göttingen, Germany; Department of Earth and Environmental Sciences, The University of Manchester, UK; Noble Research Institute, LLC, USA

## Abstract

Plant economics run on carbon and nutrients instead of money. Leaf strategies aboveground span an economic spectrum from ‘live fast and die young’ to ‘slow and steady’, but the economy defined by root strategies belowground remains unclear. Here we take a holistic view of the belowground economy, and show that root-mycorrhizal collaboration can short circuit a one-dimensional economic spectrum, providing an entire space of economic possibilities. Root trait data from 1,781 species across the globe confirm a classical fast-slow ‘conservation’ gradient but show that most variation is explained by an orthogonal ‘collaboration’ gradient, ranging from ‘do-it-yourself’ resource uptake to ‘outsourcing’ of resource uptake to mycorrhizal fungi. This broadened ‘root economics space’ provides a solid foundation for predictive understanding of belowground responses to changing environmental conditions.

**One Sentence Summary:** Collaboration broadens the ‘root economics space’ ranging from ‘do-it-yourself’ resource acquisition to ‘outsourcing’ to mycorrhizal partners.

## Main text

The diversity of plant traits across the globe shapes ecosystem functioning (*1*). Seeking general patterns, ecologists have used economic theory to explain trait variation in leaves as the aboveground plant organs for resource acquisition by photosynthesis (*1*–*3*). Aboveground plant strategies thereby fall along a ‘leaf economics spectrum’ (*2*) from cheaply-constructed but short-lived leaves optimized for ‘fast’ resource acquisition to more expensive but persistent leaves with a ‘slower’ rate of return over longer time scale.

As the belowground equivalent of leaves, fine roots acquire resources from the soil (*4*). Therefore, fine root trait variation has been hypothesized to follow a similar one-dimensional spectrum (*1*, *5*). At one side of this spectrum, plants with a ‘fast’ belowground resource acquisition strategy are expected to construct long, narrow-diameter roots with minimal biomass investment but high metabolic rates (*1*, *4*, *6*). At the opposite side of the spectrum, plants with a ‘slow’ strategy are expected to achieve longer lifespan and prolonged return on investment by constructing thicker-diameter, denser roots (*4*, *7*).

However, mixed empirical results caused ecologists to question whether variation in root traits can be adequately explained by a one-dimensional ‘fast-slow’ economics spectrum (*1*, *5*, *8*–*12*). Here, we aim to settle this debate by presenting a new conceptual framework of root economics that better captures the complexity of belowground resource acquisition strategies. First, we integrated existing knowledge to build a conceptual understanding of the covariation among four key root traits (Table 1, Fig. 1). Second, we tested our conceptual model against root traits of 1,781 plant species across all biomes of the world. All analyses were phylogenetically informed using fine-root trait data from the Global Root Trait database (GRooT) (*13*).

**Table 1.**
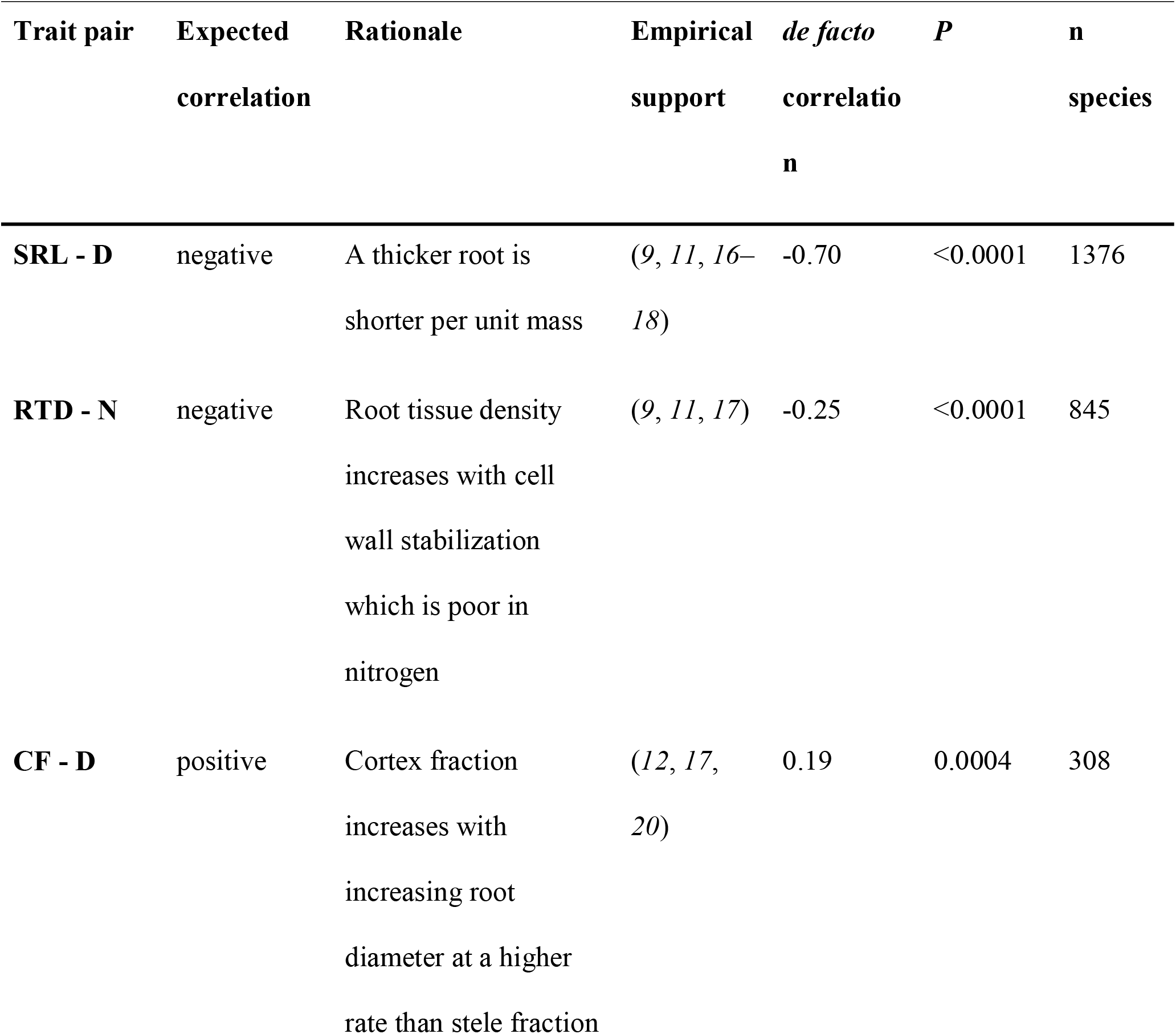
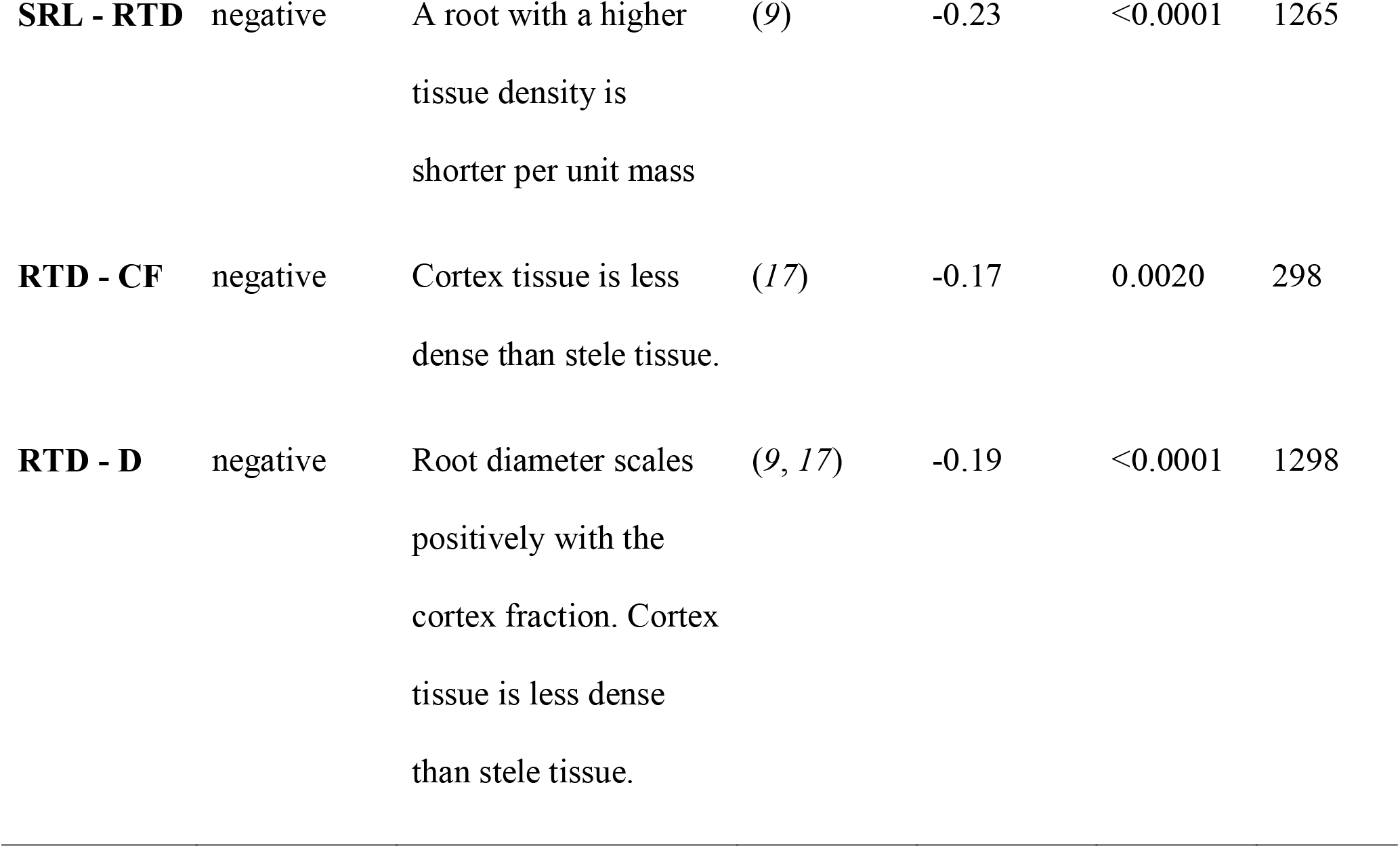
Rationale of the conceptual framework of root trait correlations depicted in Fig. 1. Expected correlations are based on mathematical and ecological rationale and empirical support from the literature. *de facto* correlations (see also fig. S1) are phylogenetically-informed correlation coefficients of species subsets with the respective trait coverage. D – root diameter, SRL – specific root length, RTD – root tissue density, N – root nitrogen content, CF - cortex fraction.

**Fig. 1.**
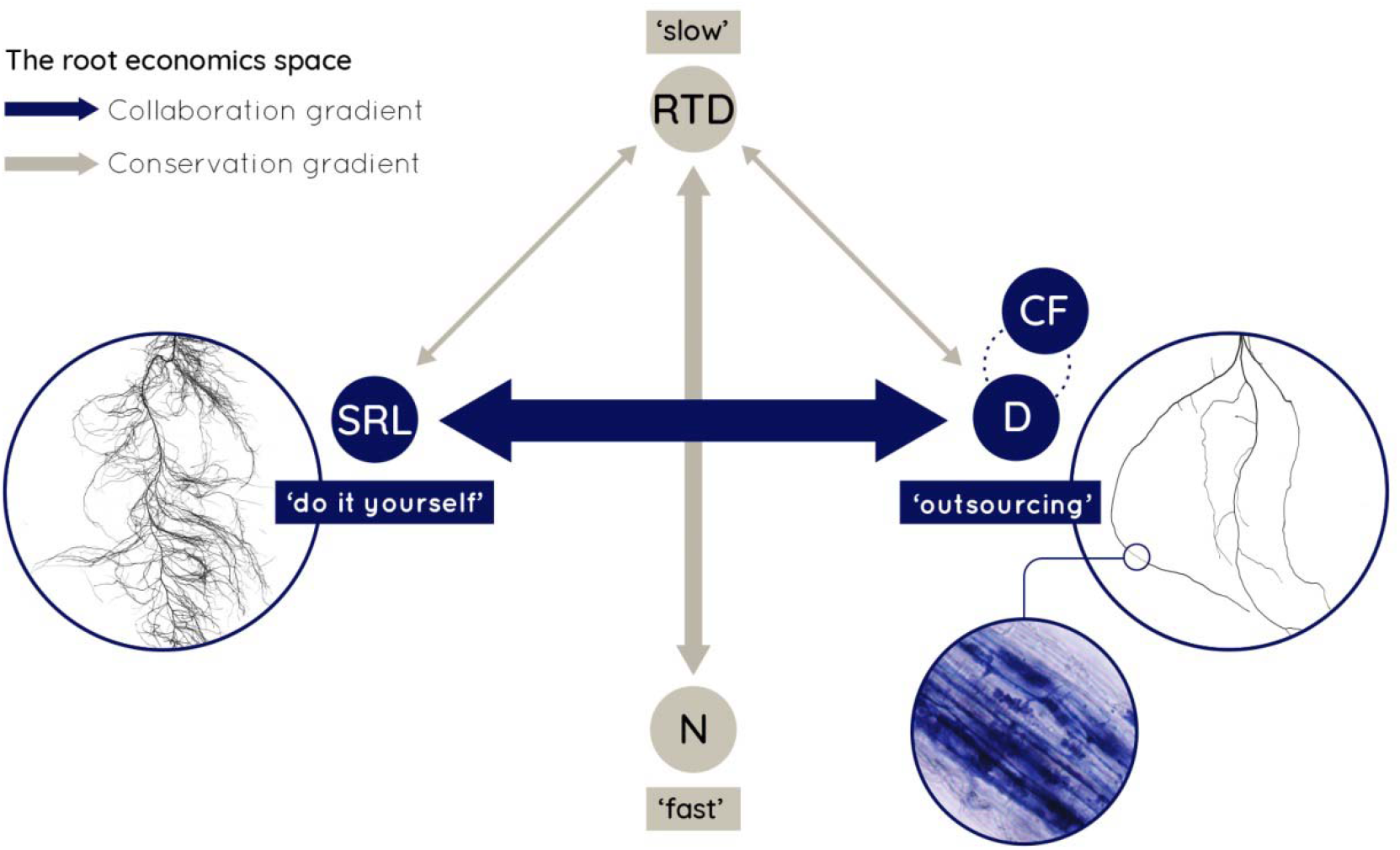
Conceptual framework of the root economics space. Based on this concept we hypothesize 1) a collaboration gradient ranging from ‘do-it-yourself’ soil exploration by high specific root length (SRL) to ‘outsourcing’ by investing carbon into the mycorrhizal partner and hence extraradical hypheae which requires a large cortex fraction (CF) and root diameter (D) and 2) a conservation gradient ranging from roots with high root tissue density (RTD) that show a ‘slow’ resource return on investment but are long-lived and well-protected, to ‘fast’ roots with a high nitrogen content (N) and metabolic rate for fast resource return on investment, but a short lifespan. Arrows indicate negative correlations between the single traits (see Table 1).

The currency of root economics is the carbon input required to construct fine roots that explore the soil for resource acquisition. Specific root length (SRL) - the root length per unit mass - therefore reflects the rate of return per unit of investment, and is a function of both root diameter (D) and root tissue density (RTD) – the root mass per unit of root volume -, following:

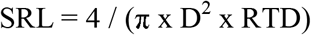

Although this equation(*6*) is a simplification when sampling heterogeneous fine root populations (*14*), it implies that SRL increases with decreasing D and/or RTD. Besides efficient soil exploration, plants have to maintain a high metabolic rate to assure ‘fast’ resource acquisition leading to high nitrogen (N) content in the fine roots (*1*, *15*). While strong negative relationships between SRL and D (*9*, *11*, *16*–*18*) as well as between RTD and N (*9*, *11*, *17*) have been observed, the relationships between SRL and RTD (*17*, *19*, *20*) as well as between D and N (*12*) have been less clear. In fact, observations across a wide range of species suggest that plants can construct roots with many combinations of SRL and RTD (*9*, *11*) indicating complex trait interactions inconsistent with a one-dimensional root economics spectrum (*8*–*12*).

We hypothesize that this root trait complexity results from the range of belowground resource uptake strategies. In contrast to aboveground photosynthesis, which is solely conducted by plant organs, belowground many species have the ability to outsource resource acquisition. This gradient of plant collaboration strategies ranges from ‘do-it-yourself’ acquisition by cheap roots for efficient soil exploration to ‘outsourcing’ acquisition via the investment of carbon in a mycorrhizal partner for the return of limiting resources. However, such outsourcing strategies have consequences for root traits. This is particularly true for arbuscular mycorrhizal fungi (AMF) because plants must increase their root cortical area, and hence their root diameter (D), to provide the intraradical habitat for their fungal partner (*17*, *21*, *22*). This is generalizable for plant symbiosis with AMF, the most widespread type of mycorrhizal fungi (*22*) and also well documented for ectomycorrhizal (EM) fungi (*23*). Thus, we hypothesize that plants can optimize resource uptake by investing carbon either in thin roots that efficiently explore the soil themselves (*9*) or in a mycorrhizal partner which requires a thick root for efficient symbiosis (Fig. 1).

This hypothesized collaboration gradient from ‘do-it-yourself’ to ‘outsourcing’ challenges the traditional spectrum of root economics that assumes D to increase with RTD for tissue conservation. Both scaling laws and empirical data (*20*) show that as D increases, root cortex area increases at a faster rate than stele area such that D scales positively with the cortex fraction (CF) (*17*) (though patterns can vary between growth forms (*12*)). The parenchymatous cortical tissue has a lower carbon content and dry weight than the stele tissue, which transports nutrients and water through lignified cells (*24*, *25*). Thus CF and RTD will be negatively correlated (Table 1). Furthermore, since D and CF are closely positively correlated, and increase in unison with mycorrhizal symbiosis, D should be negatively correlated with RTD. These relationships contradict the assumption of a one-dimensional root economics spectrum, where plants with a ‘slow’ strategy are expected to construct roots that are both thick and dense and advocate for a multi-dimensional space of root trait variation.

By testing pairwise correlations of all traits, we confirmed the bivariate relationships underlying our new concept of a belowground economics trait space with two main dimensions (Table 1). The strongest negative correlation was found between SRL and D (*R* = −0.70) representing the ‘collaboration’ gradient, from ‘do-it-yourself’ to ‘outsourcing‘. We also found a negative correlation between RTD and root N (*R* = −0.25) as observed in previous studies (*9*, *11*, *17*), which corresponds to a ‘conservation’ gradient, representing the traditional trade-off between ‘fast’ and ‘slow’ return on investment (Fig. 1).

On a sub-set of 737 species with complete information on the four main root traits (SRL, D, RTD, and root N) we could confirm these two distinct and largely independent gradients in a principal component analysis (PCA) where the first two axes encompass a plane with a cumulative explanatory power of 78% of all root trait variation. Henceforth, we refer to these gradients as the main dimensions of the **root economics space** (Fig. 2A). The first PCA axis (45% of total trait variation) represents a gradient from SRL to D, suggesting that our hypothesized ‘collaboration’ gradient is the main source of root trait variation. The second PCA axis, (33% of total trait variation) represents the ‘conservation’ gradient from root N to RTD (table S1).

**Fig. 2.**
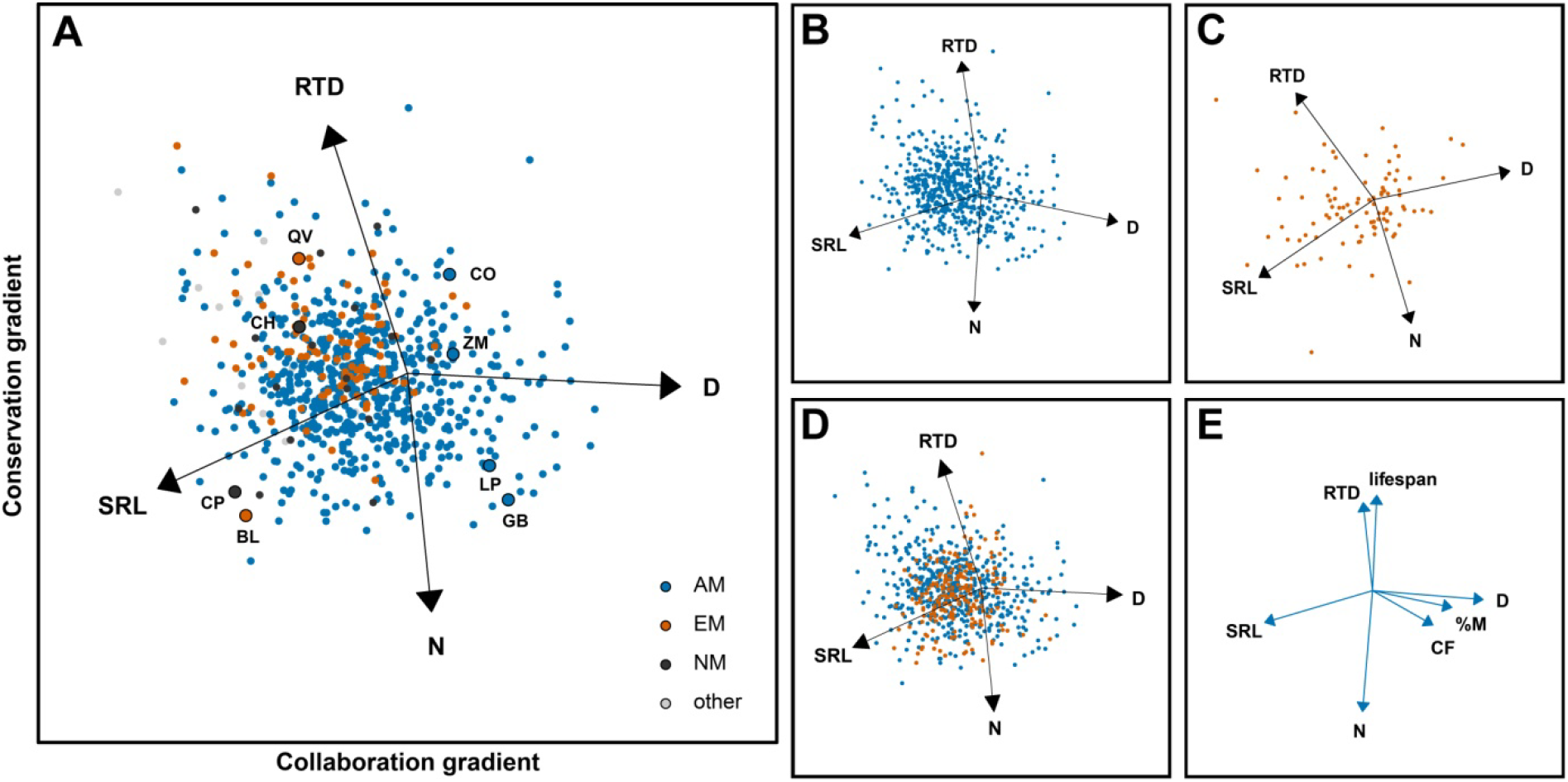
The root economics space. Phylogenetically informed principal component analyses (PCAs) of core traits of **A)** 737 species, as well as subsets of **B)** 610 arbuscular mycorrhizal (AM) species and **C)** 93 ectomycorrhizal (EM) species. The collaboration gradient (45%) ranges from ‘do-it-yourself’ roots with high specific root length (SRL) to thick diameter (D) roots with an ‘outsourcing’ strategy of nutrient acquisition. The conservation gradient (33%) explains root trait variation from ‘fast’ (high root nitrogen content– N) to ‘slow’ (high root tissue density – RTD) turnover and resource return on investment. For each corner of the **root economics space** we highlight two representative plant species: QV - *Quercus virginiana* Mill., CH - *Carex humilis* Leyss., CO - *Cornus officinalis* Siebold & Zucc., ZM - *Zea mays* L., LP - *Lathyrus pratensis* L., GB - *Ginkgo biloba* L., BL - *Betula lenta* L., CP - *Cardamine pratensis* L. **D)** Woody (blue) and non-woody (red) species show no distinct pattern within the root economics space (see also fig. S4 and table S4). **E)** PCA based on bivariate trait relationships. The percentage mycorrhizal colonization (%M) as well as the cortex fraction (CF) are positively correlated with D along the collaboration gradient, while root lifespan is negatively correlated with N along the conservation gradient. Eigenvalues, loadings and explained variances can be found in table S1. NM - non-mycorrhizal.

Species associated with AMF were the largest group in the database and were distributed over the entire trait space (Fig. 2A), but differed significantly from both non-mycorrhizal (NM) and ectomycorrhizal (EM) species (table S4). NM plants clearly aggregated on the ‘do-it-yourself’ side of the collaboration gradient, as well as on the ‘slow’ side of the conservation gradient. EM plants showed less variation along the collaboration gradient than AM plants with a tendency towards ‘do-it-yourself’ and ‘slow’ as well. A high RTD, indicative of a ‘slow’ strategy might partly originate from the fact that EM species are often woody species, although woodiness was not a significant factor of variation within the global species set (Fig. 2D, table S4). The tendency towards ‘do-it-yourself’ roots with high SRL likely results from the nature of the ectomycorrhizal symbiosis that is less dependent on cortex area but also from its more recent evolution, as evolutionarily younger species tend to have thinner roots (*9*, *21*, *25*, *26*). Even so, PCAs that solely represent the root traits of either AM or EM plant species (Fig. 2, B and C, table S1) show the same dimensions of variation as in the global dataset. Plants associated with N-fixing bacteria differed from the rest (table S4) by being located on the ‘fast’ side of the conservation gradient as their roots are rich in N (fig. S2A). Nevertheless, we could still confirm the collaboration gradient as the first PCA-axis within this species set (fig. S2, B and C, table S1). Furthermore, the two dimensions of the root economics space are present irrespective of biome or plant growth form (fig. S3 and S4, table S1).

To test our ecological interpretation of the proposed gradients, we added traits to the PCA that act as proxies for ecological functions (Fig. 2E, table S2). We used percent root length colonized by AMF (%M) as a proxy for the strength of the mycorrhizal symbiosis (*27*), and cortex fraction as a general proxy for the ability of a species to host mycorrhizal fungi (*17*, *28*, *29*). We found both %M and CF to be associated with the ‘outsourcing’ side of the collaboration gradient. To test whether the proposed conservation gradient aligns with the classical ‘fast-slow’ economics spectrum, we used root lifespan as a proxy for short- or long-term investment of plant carbon (*1*, *30*–*32*). We found that longer lifespan was indeed associated with the ‘slow’ side of the conservation gradient which is consistent with reports of negative relationships between root lifespan and N (*1*, *30*, *32*).

The decrease in root diameter over evolutionary time (*9*, *26*) suggests a reduced dependence of plants on mycorrhizal fungi. We found that the ‘collaboration’ gradient was indeed phylogenetically conserved, showing an evolutionary transition from ‘outsourcing’ to ‘do-it-yourself’ (Fig. 3, table S3 and S5). In contrast, the ‘fast-slow’ trade-off of the ‘conservation’ gradient was less pronounced across all plant families in our database (Fig. 3), and also less phylogenetically conserved (table S3). This suggests that evolutionary history causes the ‘collaboration’ gradient to be the main source of variation in root traits.

**Fig. 3.**
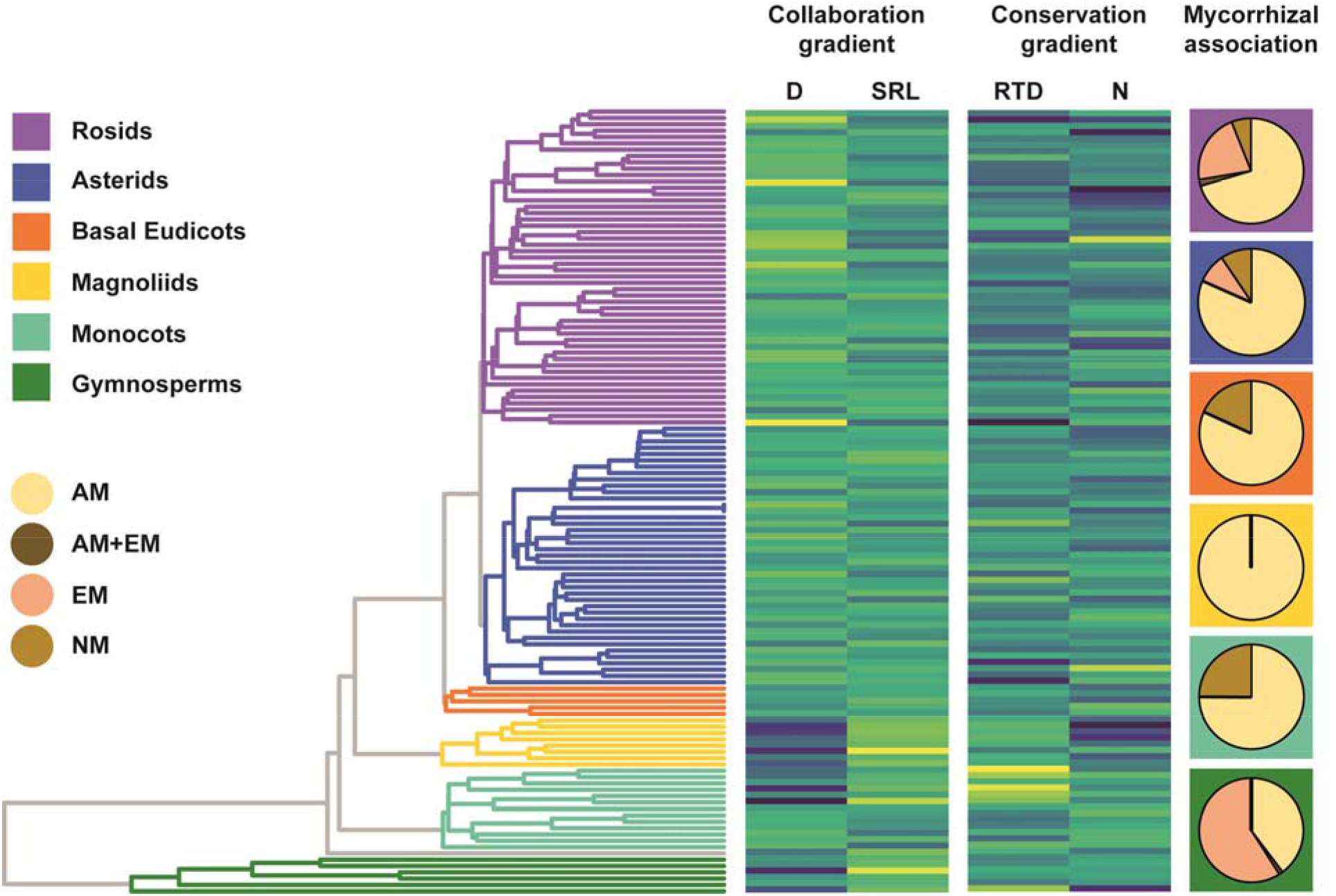
The collaboration gradient is phylogenetically conserved. Displayed is the phylogenetic tree of 1,781 species aggregated on a family level (left) with the standardized family mean trait values of the four core traits (center) ranging from low (yellow) to high (blue). The collaboration gradient shows a strong phylogenetic pattern (lambda = 0.8) with a transition from families with thick root diameter (D) to those with a high specific root length (SRL). The phylogenetic signal in the conservation gradient is less pronounced (lambda = 0.5). Pie charts (right) depict the fraction of different mycorrhizal association types within the broader plant phylogenetic clades (indicated by corresponding background colors). RTD – root tissue density, N – root nitrogen content, AM – arbuscular mycorrhizal, EM – ectomycorrhizal, NM - non-mycorrhizal.

Taken together, our results provide an answer as to why root trait variation cannot be adequately explained by a one-dimensional root economics spectrum (*8*–*11*, *17*, *33*). Plant outsourcing of belowground resource acquisition through collaboration with mycorrhizal partners represents a main dimension of root trait variation, and is fundamentally different from aboveground. This collaboration gradient from ‘do-it-yourself’ to ‘outsourcing’ represents an investment in soil exploration by either the root itself or its mycorrhizal partners. It is independent from the conservation gradient, which represents the well-known concept of ‘fast’ versus ‘slow’ return on investment. Thus both gradients depict different facets of root economics, and rather than a single one-dimensional spectrum, encompass a whole root economics space of plant strategies for belowground resource acquisition.

## Acknowledgments

We like to thank Dr. India Mansour for text editing.

## Author contributions

JB, AW and LM conceived the idea for the project; all authors were involved in collecting datasets, developing the conceptual framework and interpreting the results; JB, FvdP, DL, NG-R, OVB, and LMY performed the statistical analyses; TK annotated the mycorrhizal associations; JB, AW, CI, and LM wrote the first draft of the manuscript; all authors commented on and agreed with the final version of the manuscript. There are no conflicts of interest to declare.

## Data availability

All data analyzed in the study originate from the GRooT database(*13*) which will be publicly available at time of publication. The R script including all analyses and figure preparations is available from the corresponding author upon reasonable request.

## Supplementary Materials

Materials and Methods

Figures S1-S4

Tables S1-S5

