## Supplementary Materials for "The fungal collaboration gradient dominates the root economics space in plants"

**This PDF file includes:**

Materials and Methods
Figs. S1 to S4
Tables S1 to S5

### Materials and Methods

#### Database

All analyses presented here are based on the Global Root Trait database (GRooT) (*13*). The GRooT database combines root trait observations from FRED (*34*) and TRY (*35*) with additional datasets providing data measured on individual plants for which taxonomical information is available. It includes data on both coarse and fine roots. For the objective of this study, we selected fine roots only, as coarse roots are usually not absorptive and therefore less relevant in the context of root economics (*34*, *36*). We treated roots as fine roots if they met at least one of the following criteria: i) they were of root orders 1-3, ii) they were classified as "fine roots" by the initial authors, or iii) their diameter was smaller than 2 mm. Data measured on dead roots were excluded from the analyses. Furthermore, we excluded ferns (Polypodiopsida) because of their very special root morphology that is hardly comparable with vascular plants (*37*, *38*). We only selected data where species-level information was available. During the last decade a set of root traits was found to be highly informative of root economics: specific root length (SRL), root diameter (D), root tissue density (RTD) and root nitrogen concentration (N) (*8*–*12*, *18*, *25*). Hence, we focused our main analyses on those four traits. In addition, we analyzed the percentage of root length colonized by arbuscular mycorrhizal fungi (%M) and the root area occupied by the cortex, i.e., root cortex fraction (CF) as proxies for the strength of mycorrhizal symbiosis, as well as mean root lifespan. We checked the values of these traits for outliers, and excluded values of RTD exceeding 1.0 in further analyses.
Categorical data from GRooT such as main biome type (tropical, temperate, continental, arid or polar) following the Köppen-Geiger classification, plant woodiness (woody, non-woody or facultatively woody), mycorrhizal association (non-mycorrhizal, arbuscular mycorrhiza, ectomycorrhiza or other (e.g. ericoid mycorrhiza) and nitrogen fixing ability (fixers or non-fixers) were used in the downstream analysis and testing of our conceptual framework. GRooT includes mycorrhizal association data from FungalRoot (*39*) which did not cover our entire species set. To achieve full data cover we filled the gaps and did minor annotations based on the following general rules:
1) Mycorrhizal association is constant within species hence excluding a facultative mycorrhizal type. In cases where a lack of mycorrhizal colonization has been reported only under specific environmental conditions less suitable for the mycorrhizal symbiosis we assigned the species as mycorrhizal while in cases with intraradical hyphae but no evidence for a symbiotic interface we assigned species to be non-mycorrhizal.
2) Almost all plants have one type of mycorrhizal association as the dominant one. Therefore, dual mycorrhizal association was only assigned if species show no clear dominance towards one type.
3) The mycorrhizal association type is usually constant within a monophyletic genus and often within a family (*22*, *39*, *40*). Therefore we filled remaining gaps with the respective mycorrhizal association type of sister species.

Data processing

All data processing and analyses were done using R 3.6.1 (*41*). In this study, we analyzed how different root traits are related to each other at the level of plant species, hence a first step was to calculate species mean values. As root traits were measured in different studies varying in design (e.g. on in situ grown plants versus plants growing in pots), and because most traits varied several orders of magnitude, several steps of data processing were required before calculating species mean trait values. First, to obtain normal distributions, we log-transformed each trait, except for %M and CF, which were scaled to the range of 0-1 and arcsine square root transformed. We then *Z* transformed each trait to a mean of 0 and a standard deviation of 1 to assure variance homogeneity. Furthermore, we corrected for main study design (measurements on plants in situ, in pots or hydroponics) and the publication in which the trait measurements were first reported (as a proxy for other study specific factors, e.g. plant age, soil conditions or sample handling). This was done by building a linear mixed model for each trait, where the trait as the response variable, study design was treated as a fixed factor and publication as random factor. We used residuals of these models in further analyses.
Within some species, the categorical traits woodiness and biome had different data entries (e.g. because the species occurs both in temperate and continental biomes). In those cases, we categorized the species in the biome in which it had most observations, and we categorized its woodiness by its most commonly observed entry in further analyses.
In total, we analyzed information of 1519, 1635, 1344, and 1145 species for D, SRL, RTD and N, respectively. Scientific names in GRooT are standardized among data sets and brought up to date by querying species names using the Taxonomic Name Resolution Service v4.0 (http://tnrs.iplantcollaborative.org/) (*42*). We constructed a phylogenetic tree including all species using the backbone phylogeny from Zanne et al (*43*) and adding additional missing species with the function 'add.tips' from the package 'phangorn' (*44*). We calculated Pagel's lambda using the package 'picante' and evaluated the strength of the phylogenetic signal for each trait; a large (close to the upper bound of 1.0) Pagel's lambda value indicates higher phylogenetic conservatism (*45*), whereas a low (close to 0.0) value indicates a lack of phylogenetic conservatism.

Analyses

As all traits exerted strong phylogenetic signal (table S3), we used phylogenetically informed methods for all analyses. We first assessed bivariate relationships between the four core traits (D, SRL, RTD, N) as well as CF to build our conceptual framework (Table 1, Fig. 1), and we also tested for relationships of these traits with %M and root lifespan (fig. S1). Sample sizes varied for these bivariate correlations, depending on the number of species with complete information for both involved traits, and ranged from 19 (for the correlation between %M and root lifespan) to 1,376 (for the correlation between root diameter and specific root length) (fig. S1). In total, we used 1,781 species for these bivariate correlations. We fit Phylogenetic Generalized Least Square models using the 'pgls' function in the R package 'caper' (*46*, *47*) to each pair of traits to conduct phylogenetically corrected regression analyses. Phylogenetically corrected correlation coefficients (*r* values) were then calculated by taking the square root of the adjusted model $r^2$, and by multiplying this with -1 if the regression coefficient was negative. In cases where the adjusted model $r^2$ was negative, we assigned an *r* coefficient of 0.

We used phylogenetically informed Principal Component Analysis (PCA) to identify main dimensions of variation among root economic traits. A phylogenetic PCA was performed for the four core traits D, SRL, RTD and N, using the ‘phyl.pca’ function of the ‘phytools’ package (*48*). There were 737 species that had complete data for these 4 core traits. The eigenanalysis utilizes the correlation structure of the phylogeny to inform its estimates of eigenvalues and eigenvectors (*48*, *49*). To assess whether the PCA results and hence the dimensions of the root economics space were sensitive to biome type, mycorrhizal association, woodiness or nitrogen fixing ability, we repeated the above analysis for subsets of different biomes (tropics, temperate, continental and arid - the polar biome was represented by too few species (n=5) to perform a reliable analysis), mycorrhizal association type (arbuscular mycorrhiza versus ectomycorrhiza), woodiness (woody versus non-woody) and nitrogen fixing ability (present or absent).

We assessed whether roots from species with different mycorrhizal associations (arbuscular mycorrhiza, ectomycorrhiza, arbuscular mycorrhiza *and* ectomycorrhiza - i.e. intraspecific variation in mycorrhizal association type, ericoid mycorrhiza and non-mycorrhiza associated), species from different biomes (temperate, tropical, arid and continental), woody or non-woody species and species that either did or did not associate with bacteria able to fix nitrogen, differed significantly from each other in the global multidimensional PCA space, i.e. in their first two PCA axes (which jointly explained 77% of all trait variation). This was done using a Permutational Multiple Analysis of Variance (PERMANOVA), in which the first two PCA axes were treated as the response variables and mycorrhizal association type, biome, woodiness or ability to fix nitrogen as the fixed factor. We used Euclidean pairwise distances in PCA space among species, and calculated 999 permutations, using the ‘pairwise.adonis’ function in the ‘pairwiseAdonis’ package (*50*). To test for the significance of differences between different categories of mycorrhizal associations and biomes, we used false discovery rates (*51*) to reduce the likelihood of type I errors due to multiple testing.

Furthermore, we investigated multivariate trait space for 7 traits, i.e. the 4 core traits from the above described PCAs (D, SRL, RTD and N), supplemented by three additional traits: CF, %M and root lifespan. As PCA requires each replicate (species) to have complete data for all traits and there were few species that met this criterion, we performed an alternative dimensionality reduction analysis, based on pairwise correlations between traits. For this analysis, we used the phylogenetically informed pairwise correlations between each of the 21 trait combinations (fig. S1). We then performed a standard principal component analysis on this matrix of phylogenetic correlation coefficients.

Visual examination of the distribution of traits across the phylogeny was obtained using the function ‘phylo.heatmap’ in the package ‘phytools’ (*48*). We further examined the phylogenetic trends observed across broader phylogenetic clades of seed plants employing a randomization test to quantitatively compare individual clade trait values to the rest of the phylogeny. The test determines if the mean trait value observed in a clade deviates significantly from the population mean under the null hypothesis that the trait has a random phylogenetic distribution. To do so, we created an algorithm in R that selected clades sequentially at each node. Due to the large number of species, we selected particular nodes that enveloped important phylogenetic clades with at least 30 species (tree tips) included. For each clade we calculated the observed mean and kurtosis values as measures of central tendency and dispersion values within clades respectively. Then we generated a series of 999 random values shuffling trait values among the tips of the original tree. Significance was calculated after estimating if the observed clade mean or kurtosis were outside the 95% confidence intervals of the clade estimations using the randomized datasets. In the case of the kurtosis, values higher than the randomized mean were interpreted as evidence of underdispersion in the clade (leptokurtic distribution), whereas lower values were considered sign of overdispersion (platykurtic distribution).

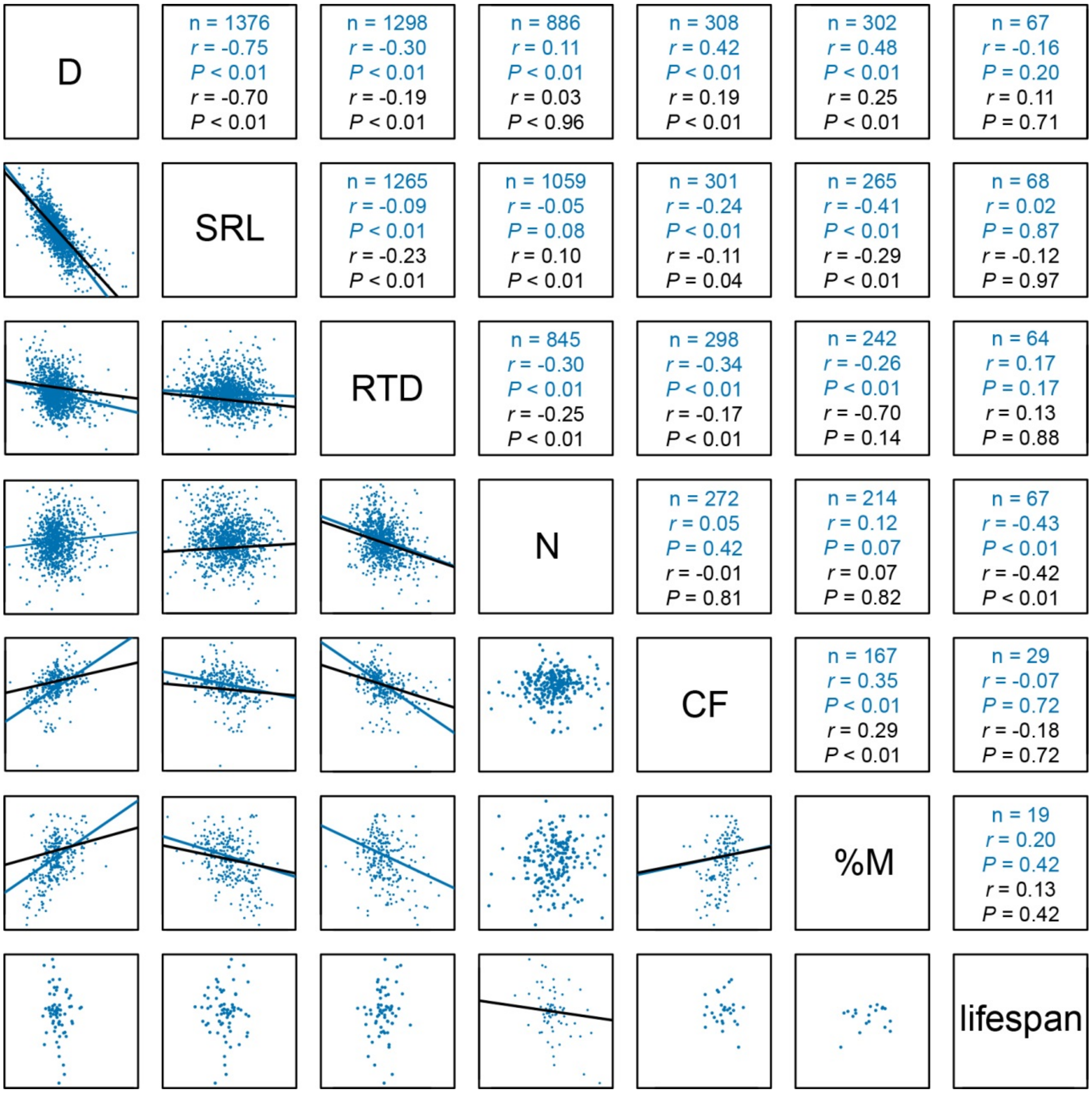


**Fig. S1. Pairwise correlation of all traits used in the analysis.** Scatterplots represent species mean trait correlations after correction for study design and publication. D – average root diameter, SRL – specific root length, RTD – root tissue density, N – root nitrogen content, CF – root cortex fraction, %M – arbuscular mycorrrhizal colonization intensity, lifespan – mean root lifespan. Regression lines represent significant correlations (blue) and significant phylogenetically corrected bivariate relationships calculated by fitting Phylogenetic Generalized Least Square models (black). Correlation coefficients are presented for the data without (blue) and with phylogenetic correction (black). Note that for the pairwise correlation of CF~%M as well as N~lifespan the two regression lines are too close to be distinguishable by eye.

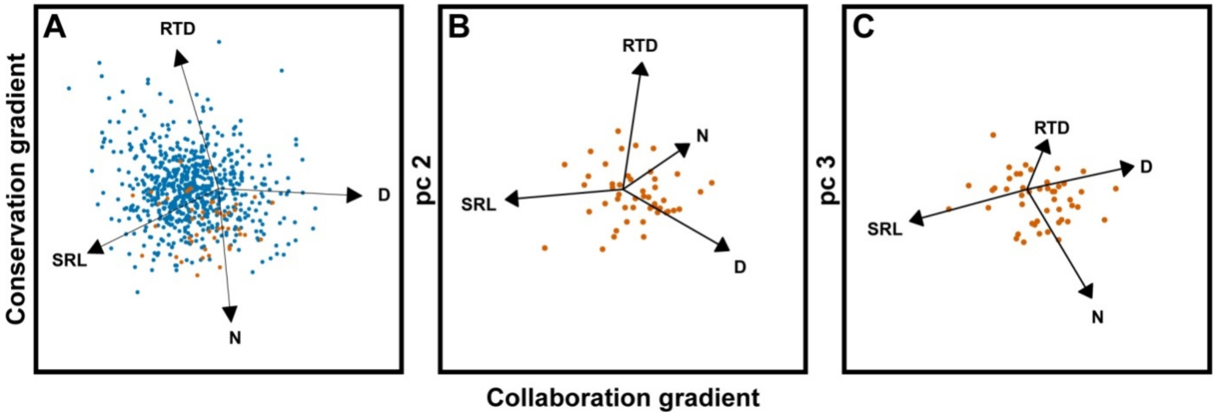


**Fig. S2. The collaboration gradient represents the main axis in the root economics space of plants associated with nitrogen fixing bacteria.** D – average root diameter, SRL – specific root length, RTD – root tissue density and N – root nitrogen content. Phylogenetically informed principal component analysis of the core species set (n=737) with plants associated with N-fixing bacteria highlighted in red (**A**). N-fixers differed from the rest by being located on the “fast” side of the conservation gradient associated with high root nitrogen content (table S4). Within the N-fixing subset the collaboration gradient appeared to be the first axis while the conservation gradient loaded on principal component 2 and 3 (**B,C**). See Table S1 for the principal component analysis.

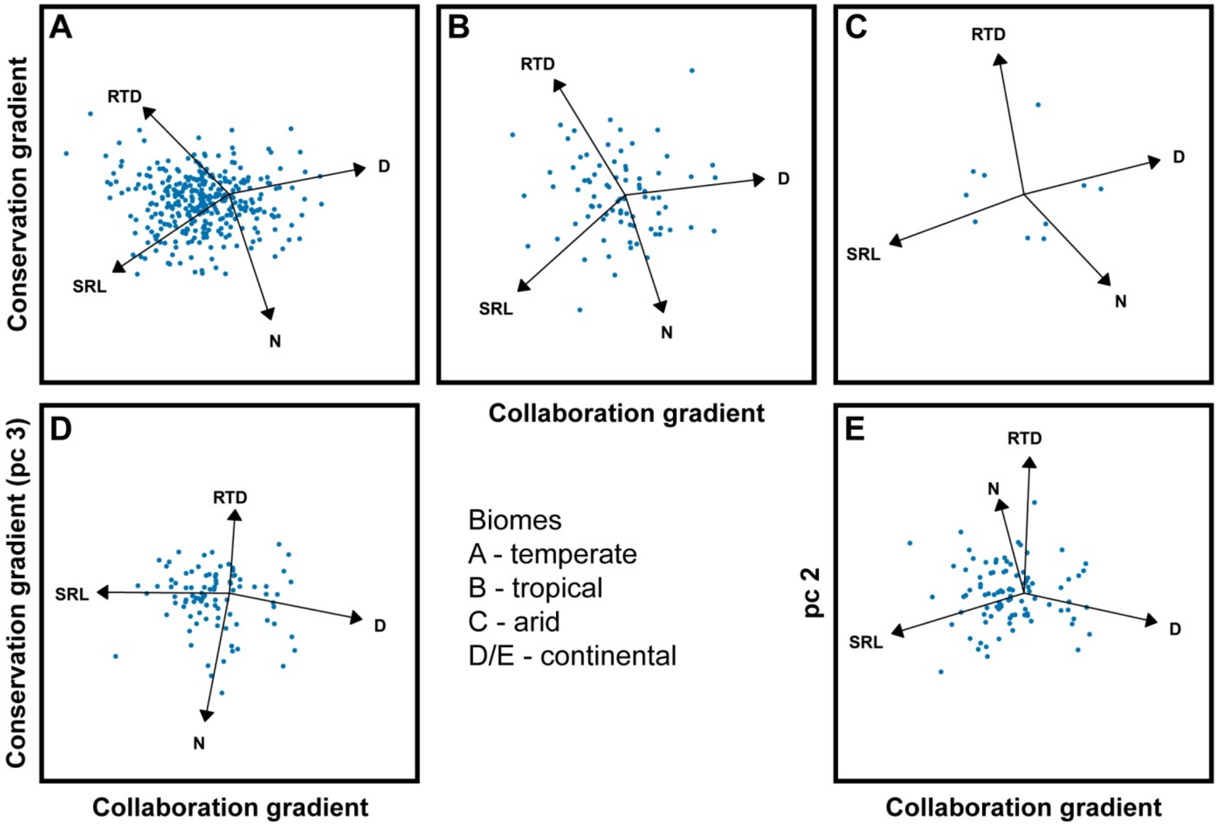


**Fig. S3. The root economics space is present in different biomes.** D – average root diameter, SRL – specific root length, RTD – root tissue density and N – root nitrogen content. Root traits and trait relations are known to vary across biomes (*9*). We found no respective between group variation within the root economics space (table S4). Still, to test whether the concept is broadly generalizable, we present separate PCAs for biomes spanning arid to tropical. We found that the root economics space was apparent in all of the biomes represented by our species. In continental systems the conservation gradient was represented by principal component 3. See table S1 for principal component analyses.

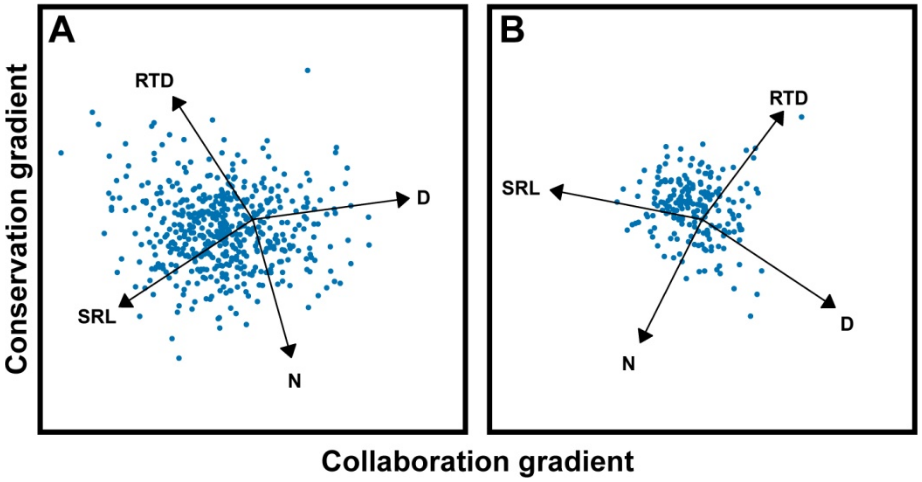


**Fig. S4. The root economics space is present in different growth forms.** D – average root diameter, SRL – specific root length, RTD – root tissue density and N – root nitrogen content. Root traits and trait relations are known to vary among plant growth forms (*12*, *25*, *52*). We found no respective between group variation within the root economics space (table S4). Still, to test whether the concept is broadly generalizable, we present separate PCAs for woody (**A**) and non-woody (**B**) species. We found that the root economics space was apparent within both plant growth forms. See table S1 for principal component analysis.

**Table S1. The root economics space can be detected irrespective of mycorrhizal association, N-fixing ability, biome or growth form.** Analysis of the core species set with full information for the four core traits. Presented here are the phylogenetically informed principal component analyses of the global species set as well as different species subsets as shown in Fig. 2A, B and C, fig S2, 3 and 4. Displayed is the Eigenvalue as well as the proportion of variance explained by each principal component (PC) and the loadings of the root traits. D – average root diameter, SRL – specific root length, RTD – root tissue density and N – root nitrogen content.

| | | PC1 | PC2 | PC3 | PC4 |
|---|---|---|---|---|---|
| **Global species** | Eigenvalue | 1.337 | 1.146 | 0.870 | 0.377 |
| n=737 | Variance | 0.447 | 0.328 | 0.189 | 0.036 |
| Fig. 2A | D | 0.963 | 0.041 | 0.062 | -0.260 |
| | SRL | -0.880 | 0.366 | 0.177 | -0.246 |
| | RTD | -0.279 | -0.784 | -0.542 | -0.120 |
| | N | 0.084 | 0.751 | -0.655 | 0.009 |
| **AM** | Eigenvalue | 1.328 | 1.149 | 0.881 | 0.373 |
| n=610 | Variance | 0.441 | 0.330 | 0.194 | 0.035 |
| Fig. 2B | D | 0.950 | 0.177 | -0.050 | 0.254 |
| | SRL | -0.917 | 0.264 | -0.170 | 0.246 |
| | RTD | -0.137 | -0.823 | 0.538 | 0.120 |
| | N | -0.052 | 0.736 | 0.675 | -0.015 |
| **EM** | Eigenvalue | 1.375 | 1.107 | 0.867 | 0.362 |
| n=93 | Variance | 0.473 | 0.306 | 0.188 | 0.033 |
| Fig. 2C | D | 0.944 | 0.173 | -0.120 | 0.252 |
| | SRL | -0.800 | -0.472 | -0.299 | 0.218 |
| | RTD | -0.542 | 0.644 | 0.523 | 0.137 |
| | N | 0.257 | -0.747 | 0.611 | 0.039 |
| **N-fixers** | Eigenvalue | 1.386 | 1.075 | 0.904 | 0.324 |
| n=48 | Variance | 0.480 | 0.289 | 0.204 | 0.026 |
| Fig. S2 | D | 0.856 | 0.443 | -0.167 | 0.208 |
| | SRL | -0.945 | 0.074 | 0.231 | 0.221 |
| | RTD | -0.154 | -0.923 | -0.336 | 0.109 |
| | N | 0.522 | -0.321 | 0.789 | 0.026 |
| **Temperate** | Eigenvalue | 1.407 | 1.082 | 0.835 | 0.388 |
| n=329 | Variance | 0.495 | 0.293 | 0.174 | 0.037 |
| Fig. S3A | D | 0.946 | -0.164 | 0.006 | 0.281 |
| | SRL | -0.807 | 0.488 | -0.228 | 0.244 |
| | RTD | -0.592 | -0.540 | 0.588 | 0.108 |
| | N | 0.291 | 0.784 | 0.547 | -0.018 |
| **Tropical** | Eigenvalue | 1.342 | 1.195 | 0.821 | 0.313 |
| n=81 | Variance | 0.450 | 0.357 | 0.168 | 0.025 |
| Fig. S3B | D | 0.967 | -0.102 | 0.099 | -0.212 |
| | SRL | -0.745 | 0.602 | 0.214 | -0.194 |
| | RTD | -0.490 | -0.722 | -0.472 | -0.130 |
| | N | 0.265 | 0.731 | -0.629 | -0.001 |

| | | | | | |
|---|---|---|---|---|---|
| **Arid** | Eigenvalue | 1.464 | 1.110 | 0.754 | 0.233 |
| n=10 | Variance | 0.536 | 0.308 | 0.142 | 0.014 |
| Fig. S3C | D | 0.943 | 0.215 | -0.195 | 0.161 |
| | SRL | -0.933 | -0.310 | 0.082 | 0.165 |
| | RTD | -0.178 | 0.877 | 0.445 | 0.026 |
| | N | 0.594 | -0.566 | 0.571 | 0.011 |
| **Continental** | Eigenvalue | 1.310 | 1.071 | 0.962 | 0.461 |
| n=94 | Variance | 0.429 | 0.287 | 0.231 | 0.053 |
| Fig. S3D/E | D | 0.919 | -0.181 | 0.163 | -0.309 |
| | SRL | -0.916 | -0.252 | -0.007 | -0.311 |
| | RTD | 0.039 | 0.848 | -0.510 | -0.139 |
| | N | -0.170 | 0.575 | 0.799 | -0.028 |
| **Woody** | Eigenvalue | 1.373 | 1.129 | 0.844 | 0.357 |
| n=517 | Variance | 0.471 | 0.319 | 0.178 | 0.032 |
| Fig. S4A | D | 0.959 | -0.117 | -0.054 | 0.252 |
| | SRL | -0.820 | 0.482 | -0.210 | 0.226 |
| | RTD | -0.487 | -0.675 | 0.543 | 0.105 |
| | N | 0.236 | 0.758 | 0.609 | -0.002 |
| **Non-woody** | Eigenvalue | 1.392 | 1.034 | 0.899 | 0.432 |
| n=209 | Variance | 0.484 | 0.267 | 0.202 | 0.047 |
| Fig. S4B | D | 0.819 | 0.486 | -0.126 | -0.277 |
| | SRL | -0.935 | -0.162 | -0.035 | -0.312 |
| | RTD | 0.496 | -0.592 | 0.626 | -0.106 |
| | N | -0.380 | 0.675 | 0.632 | 0.032 |

**Table S2. Principal component analysis based on phylogenetically corrected bivariate relationships of all trait pairs.** Displayed are the loadings of the root traits for each principal component (PC). D – average root diameter, SRL – specific root length, RTD – root tissue density, N – root nitrogen content, CF – root cortex fraction, %M – arbuscular mycorrrhizal colonization intensity, lifespan – mean root lifespan.

| | **PC1** | **PC2** | **PC3** | **PC4** | **PC5** | **PC6** | **PC7** |
|---|---|---|---|---|---|---|---|
| D | 0.603 | 0.048 | 0.113 | 0.434 | 0.119 | 0.087 | 0.641 |
| SRL | -0.588 | 0.173 | -0.373 | -0.113 | -0.050 | 0.130 | 0.673 |
| RTD | -0.050 | -0.474 | 0.528 | -0.507 | 0.112 | -0.314 | 0.354 |
| N | -0.054 | 0.660 | 0.228 | 0.056 | -0.137 | -0.697 | 0.044 |
| %C | 0.322 | 0.182 | -0.466 | -0.456 | 0.639 | -0.169 | -0.021 |
| %M | 0.426 | 0.090 | -0.204 | -0.494 | -0.714 | 0.091 | 0.083 |
| lifespan | 0.023 | -0.516 | -0.509 | 0.294 | -0.183 | -0.595 | 0.022 |

**Table S3. All traits show strong phylogenetic signal.** Displayed is Pagel´s lambda of all traits within the entire dataset as well as of the two proposed root economics gradients within the core 737 species. D – average root diameter, SRL – specific root length, RTD – root tissue density, N – root nitrogen content, CF – root cortex fraction, %M – arbuscular mycorrrhizal colonization intensity, lifespan – mean root lifespan.

| | Lambda | P |
|---|---|---|
| SRL | 0.613 | < 0.0001 |
| D | 0.784 | < 0.0001 |
| N | 0.571 | < 0.0001 |
| RTD | 0.522 | < 0.0001 |
| CF | 0.405 | < 0.0001 |
| %M | 0.586 | < 0.0001 |
| lifespan | 0.391 | 0.0005 |
| Collaboration gradient | 0.803 | < 0.0001 |
| Conservation gradient | 0.509 | < 0.0001 |

**Table S4. Permutational multivariate analysis on 737 species displaying variation between mycorrhizal types, biomes, plant growth form and nitrogen (N) fixing capacity.** AM - arbuscular mycorrhizal (n=610), EM – ectomycorrhizal (n=93), NM – non mycorrhizal (n=18), ErM – ericoid mycorrhizal (n=13), AM+EM (n=3).

| pairs | Sums of squares | *F* | $R^2$ | *P* |
|---|---|---|---|---|
| AM vs EM | 16614.13 | 24.01 | 0.033 | **0.003** |
| AM vs NM | 3992.35 | 5.48 | 0.009 | **0.020** |
| EM vs AM+EM | 1136.74 | 2.39 | 0.025 | 0.158 |
| NM vs AM+EM | 468.29 | 0.58 | 0.030 | 0.579 |
| AM vs AM+EM | 1461.42 | 2.02 | 0.003 | 0.158 |
| EM vs NM | 602.98 | 1.12 | 0.010 | 0.339 |
| AM vs ErM | 15733.96 | 21.68 | 0.034 | **0.003** |
| EM vs ErM | 5224.37 | 10.16 | 0.089 | **0.003** |
| NM vs ErM | 3203.94 | 3.85 | 0.117 | 0.060 |
| AM+EM vs ErM | 1697.42 | 2.33 | 0.143 | 0.158 |
| Temperate vs tropical | 1535.74 | 1.79 | 0.004 | 0.534 |
| Temperate vs arid | 167.14 | 0.20 | 0.001 | 0.942 |
| Tropical vs continental | 1770.56 | 2.62 | 0.015 | 0.432 |
| Tropical vs arid | 29.62 | 0.03 | 0.000 | 0.974 |
| Temperate vs continental | 914.16 | 1.25 | 0.003 | 0.573 |
| Arid vs continental | 398.04 | 0.94 | 0.009 | 0.573 |
| Woody vs non-woody | 2352.28 | 3.17 | 0.004 | 0.144 |
| N-fixing vs non-fixing | 12604.00 | 17.50 | 0.023 | **0.001** |

**Table S5. Mean and kurtosis values of four fine root functional traits for multiple clades in the Spermatophytes.** Values are standardized to represent deviation from the mean of each trait, so negative and positive values indicate clade relative mean deviation from the mean value for the entire phylogeny (1,781 species). Bold values represent clades that had mean or kurtosis values different (95% confidence interval) from the overall values for the entire phylogeny. Positive kurtosis values indicate underdispersion of values in the clade, whereas negative values represent overdispersion. 'n' indicates the number of species per clade and 'se' the standard error of the mean. Phylogenetic patterns in root trait distribution confirm the previous observations that ancestry explains a remarkable amount of variation in seed plants and contrasting trait patterns among numerous clades. For instance, our study confirms the relative large root diameter (D) prevalent in the Magnoliids and Gymnosperms compared to other seed plant groups (*52*). Magnoliids also showed lower root tissue density (RTD) and Magnoliales higher root nitrogen content (N) than other seed plants, highlighting the particular set of traits in this group. On the other hand, Ericales (Asterids), Caryophyllales and Myrtales showed smaller D, higher specific root length (SRL) and RTD than other clades. Monocots and Poales showed high SRL and low N compared to other plants. The clade of Fagales and Fabales also showed higher RTD and N than most clades, possibly showing the ubiquitous association of these species with N-fixing bacteria and the dominance of woody species (*53, 54*). Most of the Lamiidae clades showed low RTD values indicating the dominance of non-woody plants in these clades. Patterns in the evenness of the distribution among clades were also interesting. Campanuliids showed a remarkable conservatism in D, whereas Fabales and Fagales showed tight variation in N content. On the other hand, Magnoliids showed more intraspecific variation in D than expected by chance, and Fagales showed a wide range of N values. Interestingly, RTD values showed a random pattern distribution across all clades, confirming the low phylogenetic signal found in this study and previously reported for this trait (25).

**Diameter**

| Clade | n | mean | se | 95% CI (mean) | | kurtosis | 95% CI (kurtosis) | |
|---|---|---|---|---|---|---|---|---|
| Spermatophyta | 1519 | 0.00 | 0.01 | -0.03 | 0.03 | 3.16 | -1.32 | 1.23 |
| Angiosperms | 1451 | -0.01 | 0.01 | -0.03 | 0.02 | 3.22 | -1.28 | 1.34 |
| Monocots | 297 | 0.01 | 0.03 | -0.05 | 0.07 | 4.31 | -1.71 | 3.43 |
| Poales | 264 | -0.04 | 0.03 | -0.10 | 0.02 | 5.74 | -0.46 | 4.92 |
| Poaceae | 199 | -0.04 | 0.03 | -0.11 | 0.03 | 4.79 | -1.95 | 4.20 |
| Eudicots | 1154 | -0.02 | 0.02 | -0.04 | 0.02 | 2.98 | -1.71 | 1.21 |
| Core Eudicots | 1068 | **-0.07** | **0.02** | **-0.10** | **-0.04** | 4.40 | -0.33 | 2.69 |
| Asterids + Caryophyllales | 483 | 0.00 | 0.02 | -0.04 | 0.05 | 5.32 | -0.14 | 4.08 |
| Caryophyllales | 55 | **-0.19** | **0.08** | **-0.33** | **-0.05** | 0.34 | -7.48 | 0.81 |
| Asterids | 444 | 0.02 | 0.02 | -0.02 | 0.07 | 5.29 | -0.28 | 4.11 |
| Ericales + Cornales | 79 | **-0.14** | **0.07** | **-0.25** | **-0.02** | 0.39 | -7.48 | 0.54 |
| Ericales | 67 | **-0.20** | **0.06** | **-0.33** | **-0.07** | 0.43 | -7.45 | 0.72 |
| Campanuliids + Lamiids | 365 | 0.06 | 0.02 | -0.01 | 0.11 | **7.22** | **1.49** | **6.19** |
| Campanuliids | 223 | 0.10 | 0.03 | -0.04 | 0.17 | **9.10** | **2.59** | **8.39** |
| Asterales | 175 | 0.10 | 0.03 | -0.02 | 0.18 | **11.79** | **4.99** | **11.27** |
| Asteraceae | 102 | 0.09 | 0.03 | -0.01 | 0.19 | **12.12** | **4.45** | **12.08** |

| | | | | | | | | |
|---|---|---|---|---|---|---|---|---|
| Lamiids | 142 | -0.02 | 0.04 | -0.10 | 0.07 | 1.62 | -5.66 | 1.29 |
| Lamiales | 82 | 0.03 | 0.03 | -0.08 | 0.14 | 0.47 | -7.30 | 0.62 |
| Gentianales | 44 | -0.08 | 0.09 | -0.23 | 0.07 | 0.24 | -7.35 | 0.86 |
| Rosids + Saxifragales | 553 | **-0.14** | **0.02** | **-0.18** | **-0.10** | 4.19 | -1.14 | 2.88 |
| Rosids | 539 | **-0.14** | **0.02** | **-0.18** | **-0.10** | 4.24 | -1.13 | 2.93 |
| Fabids | 361 | **-0.14** | **0.02** | **-0.19** | **-0.08** | 0.80 | -4.93 | 2.25 |
| Malpighiales | 66 | **-0.33** | **0.05** | **-0.46** | **-0.21** | 0.16 | -7.62 | 0.47 |
| Fabaceae | 116 | **0.14** | **0.04** | **0.05** | **0.23** | 1.41 | -6.17 | 1.25 |
| Rosales | 91 | **-0.21** | **0.04** | **-0.31** | **-0.10** | 0.38 | -7.61 | 0.42 |
| Rosaceae | 61 | **-0.17** | **0.05** | **-0.30** | **-0.04** | 0.13 | -7.66 | 0.50 |
| Fagales | 75 | **-0.31** | **0.04** | **-0.43** | **-0.19** | 0.68 | -7.35 | 0.90 |
| Fagaceae | 49 | **-0.26** | **0.05** | **-0.40** | **-0.12** | 0.72 | -6.53 | 1.25 |
| Malvids | 174 | -0.17 | 0.05 | -0.24 | 0.09 | 5.29 | -1.70 | 4.80 |
| Malvales | 41 | 0.03 | 0.11 | -0.13 | 0.19 | 4.28 | -2.80 | 4.94 |
| Sapindales | 64 | -0.09 | 0.08 | -0.21 | 0.04 | 4.23 | -3.71 | 4.56 |
| Myrtales | 43 | **-0.45** | **0.06** | **-0.61** | **-0.30** | -0.92 | -8.31 | -0.29 |
| Myrtaceae | 31 | **-0.55** | **0.08** | **-0.74** | **-0.37** | -0.88 | -7.52 | 0.00 |
| Basal Eudicots | 32 | 0.05 | 0.06 | -0.14 | 0.22 | 0.75 | -5.90 | 1.62 |
| Magnoliids | 86 | **0.66** | **0.05** | **0.55** | **0.77** | **-0.32** | **-8.31** | **-0.25** |
| Laurales | 45 | **0.62** | **0.08** | **0.47** | **0.77** | -0.61 | -7.79 | 0.05 |
| Lauraceae | 42 | **0.58** | **0.08** | **0.43** | **0.73** | -0.25 | -7.63 | 0.44 |
| Magnoliales | 40 | **0.70** | **0.05** | **0.54** | **0.86** | -0.49 | -7.70 | 0.21 |
| Magnoliaceae | 33 | **0.71** | **0.05** | **0.53** | **0.88** | -0.62 | -7.49 | 0.22 |
| Gymnosperms | 65 | **0.15** | **0.05** | **0.02** | **0.27** | 3.77 | -4.13 | 4.09 |
| Pinales | 62 | **0.16** | **0.04** | **0.03** | **0.28** | 2.83 | -4.92 | 3.20 |
| Cupressales | 31 | **0.27** | **0.07** | **0.08** | **0.44** | 1.42 | -4.85 | 2.32 |
| Pinaceae | 31 | 0.05 | 0.04 | -0.13 | 0.23 | 0.14 | -6.38 | 1.03 |

**Specific root length**

| Clade | n | mean | se | 95% CI (mean) | | kurtosis | 95% CI (kurtosis) | |
|---|---|---|---|---|---|---|---|---|
| Spermatophyta | 1634 | 0.01 | 0.01 | -0.02 | 0.02 | 0.17 | -0.22 | 0.21 |
| Angiosperms | 1566 | 0.02 | 0.01 | -0.01 | 0.03 | 0.15 | -0.24 | 0.19 |
| Monocots | 222 | **0.16** | **0.03** | **0.09** | **0.21** | 0.28 | -0.49 | 0.69 |
| Poales | 188 | **0.21** | **0.03** | **0.13** | **0.26** | 0.42 | -0.35 | 0.85 |
| Poaceae | 158 | **0.21** | **0.03** | **0.13** | **0.28** | 0.55 | -0.28 | 1.04 |
| Eudicots | 1341 | 0.00 | 0.01 | -0.04 | 0.01 | 0.14 | -0.26 | 0.20 |
| Core Eudicots | 1220 | 0.03 | 0.01 | -0.01 | 0.04 | 0.25 | -0.16 | 0.33 |
| Asterids + Caryphyllales | 605 | 0.02 | 0.02 | -0.03 | 0.05 | 0.25 | -0.27 | 0.43 |
| Asterids | 535 | 0.01 | 0.02 | -0.04 | 0.04 | 0.29 | -0.26 | 0.50 |
| Ericales + Cornales | 81 | **0.13** | **0.06** | **0.02** | **0.22** | 0.50 | -0.61 | 1.24 |
| Campanuliids + Lamiids | 454 | -0.01 | 0.02 | -0.06 | 0.02 | 0.19 | -0.37 | 0.42 |
| Campanuliids | 294 | -0.03 | 0.02 | -0.09 | 0.01 | 0.04 | -0.63 | 0.38 |
| Asterales | 258 | -0.03 | 0.03 | -0.09 | 0.02 | 0.17 | -0.53 | 0.54 |
| Asteraceae | 217 | -0.03 | 0.03 | -0.10 | 0.02 | -0.09 | -0.84 | 0.33 |

| | | | | | | | | |
|---|---|---|---|---|---|---|---|---|
| Lamiids | 160 | 0.03 | 0.04 | -0.06 | 0.09 | 0.12 | -0.71 | 0.63 |
| Lamiales | 94 | -0.05 | 0.05 | -0.16 | 0.03 | 0.43 | -0.60 | 1.11 |
| Gentianales | 47 | **0.16** | **0.07** | **0.02** | **0.29** | -0.48 | -1.79 | 0.48 |
| Caryophyllales | 46 | **0.16** | **0.07** | **0.01** | **0.28** | -0.19 | -1.51 | 0.78 |
| Rosids + Saxifragales | 615 | 0.03 | 0.02 | -0.01 | 0.06 | 0.26 | -0.26 | 0.43 |
| Rosids | 600 | 0.04 | 0.02 | -0.01 | 0.06 | 0.26 | -0.26 | 0.45 |
| Fabidae | 408 | 0.00 | 0.02 | -0.05 | 0.04 | 0.09 | -0.50 | 0.34 |
| Malpighiales | 69 | 0.24 | 0.05 | 0.11 | 0.34 | -0.59 | -1.74 | 0.21 |
| Fabaceae | 140 | -0.26 | 0.04 | -0.34 | -0.19 | -0.25 | -1.16 | 0.28 |
| Rosales | 103 | **0.11** | **0.05** | **0.01** | **0.19** | -0.04 | -1.01 | 0.59 |
| Fagales | 76 | 0.12 | 0.04 | 0.00 | 0.21 | **1.80** | **0.69** | **2.55** |
| Fagaceae | 49 | **0.07** | **0.05** | **-0.07** | **0.19** | **3.21** | **1.93** | **4.15** |
| Malviids | 192 | **0.11** | **0.04** | **0.03** | **0.17** | 0.47 | -0.30 | 0.90 |
| Malvales | 42 | -0.08 | 0.08 | -0.23 | 0.06 | -0.43 | -1.79 | 0.59 |
| Sapindales | 67 | 0.03 | 0.06 | -0.09 | 0.14 | 0.63 | -0.54 | 1.44 |
| Magnoliids | 86 | **-0.42** | **0.03** | **-0.53** | **-0.33** | -0.09 | -1.15 | 0.62 |
| Laurales | 45 | **-0.43** | **0.05** | **-0.58** | **-0.30** | -0.44 | -1.78 | 0.53 |
| Lauracaea | 42 | **-0.42** | **0.05** | **-0.58** | **-0.28** | -0.50 | -1.93 | 0.53 |
| Magnoliales | 40 | **-0.40** | **0.04** | **-0.55** | **-0.26** | 0.23 | -1.21 | 1.26 |
| Magnoliaceae | 34 | **-0.42** | **0.04** | **-0.59** | **-0.28** | 0.45 | -0.99 | 1.55 |
| Gymnosperms | 68 | **-0.23** | **0.04** | **-0.35** | **-0.13** | 5.32 | 4.16 | 6.13 |
| Pinales | 65 | **-0.24** | **0.04** | **-0.37** | **-0.14** | 0.34 | -0.85 | 1.17 |
| Cupressales | 31 | **-0.30** | **0.06** | **-0.48** | **-0.15** | -0.53 | -1.94 | 0.61 |
| Pinaceae | 34 | **-0.19** | **0.05** | **-0.36** | **-0.04** | 1.05 | -0.39 | 2.16 |

**Root tissue density**

| Clade | n | mean | se | 95% CI (mean) | | kurtosis | 95% CI (kurtosis) | |
|---|---|---|---|---|---|---|---|---|
| Spermatophyta | 1344 | 0.02 | 0.02 | 1.25 | -0.03 | 0.03 | -0.61 | 0.59 |
| Angiopsperms | 1292 | 0.02 | 0.02 | 1.19 | -0.03 | 0.03 | -0.71 | 0.54 |
| Monocots | 219 | 0.08 | 0.04 | 0.68 | -0.02 | 0.14 | -2.08 | 0.68 |
| Poales | 194 | **0.16** | **0.04** | **0.01** | **0.05** | 0.22 | -2.89 | 0.05 |
| Poaceae | 120 | 0.03 | 0.04 | 1.15 | -0.11 | 0.11 | -2.18 | 1.41 |
| Eudicots | 1070 | 0.01 | 0.02 | 1.36 | -0.05 | 0.02 | -0.61 | 0.75 |
| Core Eudicots | 984 | 0.04 | 0.02 | 1.36 | -0.02 | 0.06 | -0.61 | 0.79 |
| Asterids + Caryophyllales | 450 | -0.08 | 0.03 | 0.88 | -0.16 | -0.05 | -1.47 | 0.56 |
| Asterids | 420 | -0.11 | 0.03 | 1.27 | -0.19 | -0.07 | -1.12 | 0.99 |
| Ericales | 59 | 0.16 | 0.08 | 0.99 | -0.02 | 0.28 | -2.89 | 1.60 |
| Campanuliids | 215 | -0.17 | 0.03 | 1.13 | -0.28 | -0.12 | -1.69 | 1.13 |
| Asterales | 180 | -0.20 | 0.03 | 1.56 | -0.31 | -0.14 | -1.36 | 1.63 |
| Asteraceae | 148 | -0.21 | 0.03 | 1.02 | -0.34 | -0.14 | -2.10 | 1.18 |
| Lamiids | 133 | -0.13 | 0.05 | 0.63 | -0.26 | -0.06 | -2.54 | 0.82 |
| Lamiales | 87 | **-0.17** | **0.06** | **-0.17** | **-0.32** | -0.07 | -3.68 | 0.24 |
| Lamiaceae | 63 | **-0.17** | **0.07** | **-0.33** | **-0.34** | -0.04 | -4.10 | 0.23 |
| Gentianales | 42 | -0.09 | 0.09 | 1.39 | -0.29 | 0.07 | -2.50 | 2.19 |

| | | | | | | | | |
|---|---|---|---|---|---|---|---|---|
| Rosids + Saxifragales | 504 | **0.17** | **0.03** | **1.14** | **0.09** | 0.20 | -1.10 | 0.81 |
| Rosids | 490 | **0.17** | **0.03** | **1.13** | **0.09** | 0.20 | -1.14 | 0.80 |
| Fabiids | 322 | **0.15** | **0.03** | **1.15** | **0.06** | 0.19 | -1.38 | 0.98 |
| Malpighiales | 67 | 0.05 | 0.08 | 1.28 | -0.11 | 0.17 | -2.54 | 1.84 |
| Fabales | 255 | **0.18** | **0.04** | **1.14** | **0.08** | 0.23 | -1.53 | 1.07 |
| Fabaceae | 100 | 0.02 | 0.05 | 1.52 | -0.12 | 0.11 | -1.90 | 1.85 |
| Rosales | 82 | **0.19** | **0.07** | **0.21** | **0.03** | 0.29 | -3.48 | 0.63 |
| Fagales | 68 | **0.38** | **0.08** | **1.68** | **0.22** | 0.50 | -2.02 | 2.22 |
| Fagaceae | 46 | 0.40 | 0.08 | -0.03 | 0.20 | 0.55 | -4.09 | 0.72 |
| Malviids | 166 | **0.19** | **0.05** | **0.94** | **0.08** | 0.26 | -2.10 | 1.05 |
| Malvales | 39 | 0.20 | 0.12 | -0.75 | -0.02 | 0.36 | -4.64 | 0.10 |
| Myrtales | 41 | **0.36** | **0.10** | **0.28** | **0.15** | 0.53 | -3.45 | 1.10 |
| Myrtaceae | 32 | 0.47 | 0.12 | -0.29 | 0.23 | 0.64 | -4.02 | 0.68 |
| Magnoliids | 86 | **-0.37** | **0.05** | **-0.01** | **-0.52** | -0.27 | -3.66 | 0.42 |
| Laurales | 45 | **-0.30** | **0.07** | **-0.67** | **-0.50** | -0.15 | -4.53 | 0.10 |
| Lauraceae | 40 | -0.45 | 0.06 | 1.35 | -0.66 | -0.29 | -2.64 | 2.18 |
| Magnoliales | 34 | -0.46 | 0.05 | 0.48 | -0.69 | -0.29 | -3.40 | 1.42 |
| Magnoliaceae | 33 | -0.48 | 0.05 | 0.75 | -0.71 | -0.30 | -2.97 | 1.73 |
| Gymnosperms | 52 | 0.04 | 0.05 | 0.71 | -0.15 | 0.18 | -3.28 | 1.39 |

**Root nitrogen content**

| Clade | n | mean | se | 95% CI (mean) | | kurtosis | 95% CI (kurtosis) | |
|---|---|---|---|---|---|---|---|---|
| Spermatophyta | 1144 | 0.00 | 0.02 | 0.54 | -0.04 | 0.04 | -0.32 | 0.30 |
| Angiosperms | 1089 | 0.00 | 0.02 | 0.44 | -0.03 | 0.04 | -0.42 | 0.21 |
| Monocots | 190 | **-0.33** | **0.04** | **-0.17** | **-0.42** | -0.24 | -1.49 | 0.00 |
| Poales | 172 | **-0.33** | **0.05** | **-0.22** | **-0.42** | -0.23 | -1.56 | -0.01 |
| Poaceae | 155 | **-0.30** | **0.05** | **-0.01** | **-0.39** | -0.20 | -1.37 | 0.26 |
| Eudicots | 896 | **0.07** | **0.02** | **0.49** | **0.04** | 0.12 | -0.41 | 0.29 |
| Core Eudicots | 820 | 0.04 | 0.02 | 0.57 | 0.00 | 0.09 | -0.34 | 0.38 |
| Asterids + Caryophyllales | 405 | -0.05 | 0.03 | 0.63 | -0.10 | 0.01 | -0.45 | 0.58 |
| Asterids | 352 | -0.05 | 0.03 | 0.66 | -0.11 | 0.02 | -0.44 | 0.66 |
| Ericales + Cornales | 69 | -0.30 | 0.06 | 2.14 | -0.44 | -0.16 | 0.39 | 2.75 |
| Campanuliids + Lamiids | 283 | 0.01 | 0.03 | 0.33 | -0.06 | 0.09 | -0.84 | 0.40 |
| Campanuliids | 189 | -0.03 | 0.04 | 0.34 | -0.12 | 0.06 | -0.96 | 0.52 |
| Asterales | 134 | 0.00 | 0.04 | 0.83 | -0.10 | 0.11 | -0.63 | 1.13 |
| Asteraceae | 130 | 0.00 | 0.05 | 0.87 | -0.10 | 0.10 | -0.60 | 1.16 |
| Lamiids | 94 | 0.09 | 0.06 | 0.54 | -0.03 | 0.22 | -1.08 | 1.00 |
| Lamiales | 52 | 0.01 | 0.08 | 0.45 | -0.15 | 0.18 | -1.36 | 1.19 |
| Gentianales | 34 | **0.16** | **0.08** | **-0.63** | **-0.05** | 0.37 | -2.59 | 0.36 |
| Caryophyllales | 53 | **-0.04** | **0.09** | **-0.03** | **-0.20** | 0.13 | -1.88 | 0.70 |
| Caryophyllaceae | 33 | **-0.09** | **0.13** | **-0.69** | **-0.12** | 0.30 | -2.75 | 0.33 |
| Rosids + Saxifragales | 394 | **0.14** | **0.03** | **0.15** | **0.08** | 0.20 | -0.93 | 0.11 |
| Rosids | 380 | **0.15** | **0.03** | **0.11** | **0.09** | 0.22 | -0.97 | 0.09 |
| Fabiids | 271 | **0.25** | **0.04** | **0.35** | **0.18** | **0.32** | **-1.44** | **-0.18** |

| | | | | | | | | |
|---|---|---|---|---|---|---|---|---|
| Malpighiales | 48 | 0.04 | 0.07 | 1.00 | -0.13 | 0.22 | -0.94 | 1.80 |
| Fabales | 209 | **0.30** | **0.05** | **0.62** | **0.22** | **0.39** | **1.89** | **0.28** |
| Fabaceae | 75 | **0.83** | **0.08** | **0.95** | **0.69** | **0.97** | **2.06** | **0.22** |
| Fagales + Rosales | 134 | 0.01 | 0.05 | 1.51 | -0.09 | 0.12 | 0.07 | 1.80 |
| Rosales | 72 | -0.01 | 0.08 | 0.69 | -0.15 | 0.13 | -1.01 | 1.26 |
| Rosaceae | 45 | **-0.18** | **0.06** | **-0.32** | **-0.36** | 0.00 | -2.21 | 0.51 |
| Fagales | 72 | **0.15** | **0.07** | **0.76** | **0.09** | 0.19 | -2.47 | -0.19 |
| Fagaceae | 59 | **0.22** | **0.06** | **4.93** | **0.18** | **0.14** | **3.16** | **5.61** |
| Malviids | 109 | -0.08 | 0.06 | 0.24 | -0.19 | 0.04 | -1.25 | 0.62 |
| Sapindales | 45 | **-0.04** | **0.08** | **-0.80** | **-0.22** | 0.15 | -2.74 | 0.04 |
| Magnoliids | 76 | 0.43 | 0.05 | -0.02 | 0.30 | 0.58 | -1.74 | 0.52 |
| Laurales | 38 | 0.53 | 0.07 | -0.61 | 0.34 | 0.73 | -2.53 | 0.32 |
| Lauraceae | 35 | 0.48 | 0.07 | -0.37 | 0.29 | 0.69 | -2.43 | 0.60 |
| Magnoliales | 33 | **0.31** | **0.07** | **0.19** | **0.10** | 0.52 | -1.83 | 1.21 |
| Magnolinaceae | 32 | **0.34** | **0.06** | **0.25** | **0.13** | 0.56 | -1.73 | 1.28 |
| Gymnosperms | 55 | **-0.15** | **0.04** | **-3.47** | **-0.32** | 0.01 | 1.62 | 4.19 |
| Pinales | 54 | **-0.18** | **0.03** | **-0.20** | **-0.34** | -0.01 | -2.05 | 0.54 |
| Pinaceae | 32 | **-0.13** | **0.05** | **-0.42** | **-0.34** | 0.08 | -1.53 | 1.46 |
